# mulea - an R package for enrichment analysis using multiple ontologies and empirical FDR correction

**DOI:** 10.1101/2024.02.28.582444

**Authors:** Cezary Turek, Márton Ölbei, Tamás Stirling, Gergely Fekete, Ervin Tasnádi, Leila Gul, Balázs Bohár, Balázs Papp, Wiktor Jurkowski, Eszter Ari

## Abstract

Traditional gene set enrichment analyses are typically limited to a few ontologies and do not account for the interdependence of gene sets or terms, resulting in overcorrected *p*-values. To address these challenges, we introduce *mulea*, an R package offering comprehensive overrepresentation and functional enrichment analysis. *mulea* employs an innovative *empirical false discovery rate (eFDR) correction method*, specifically designed for interconnected biological data, to accurately identify significant terms within diverse ontologies. *mulea* expands beyond traditional tools by incorporating a wide range of ontologies, encompassing Gene Ontology, pathways, regulatory elements, genomic locations, and protein domains. This flexibility enables researchers to tailor enrichment analysis to their specific questions, such as identifying enriched transcriptional regulators in gene expression data or overrepresented protein domains in protein sets. To facilitate seamless analysis, *mulea* provides gene sets (in standardised GMT format) for 27 model organisms, covering 16 databases and various identifiers resulting in almost 900 files. Additionally, the *muleaData* ExperimentData Bioconductor package simplifies access to these pre-defined ontologies. Finally, *mulea*’s architecture allows for easy integration of user-defined ontologies, expanding its applicability across diverse research areas.

**Availability and Implementation:** Software for the tools demonstrated in this article is available as an *R* package on GitHub: https://github.com/ELTEbioinformatics/mulea.

## Introduction

Large-scale omics studies, such as transcriptomic and proteomic, often generate extensive lists of genes, transcripts, or proteins exhibiting differential expression or specific characteristics. However, deciphering the biological mechanisms underlying these gene lists can be challenging. *Overrepresentation analysis* (ORA) and *gene set enrichment analysis* (GSEA) help extract meaningful insights by identifying shared characteristics among these genes, transcripts, or proteins. While widely used tools (Kuleshov *et al*., 2016; Raudvere *et al*., 2019; Wu *et al*., 2021) typically focus on Gene Ontology (GO) (The Gene Ontology Consortium, 2021) or KEGG pathway (Kanehisa *et al*., 2021) enrichment, incorporating additional gene and protein properties can offer deeper understanding. Prompted by the lack of such comprehensive approaches, we developed the R package *mulea*: multi-enrichment analysis, which enables enrichment analyses using a diverse range of gene sets and ontologies.

*mulea* enables overrepresentation testing for a wide range of ontologies, including pathways, protein domains, genomic locations, GO terms, and gene expression regulators (such as transcription factors and microRNAs). We provide these ontologies from 16 publicly available databases, in a standardised GMT (*Gene Matrix Transposed*) format (https://github.com/ELTEbioinformatics/GMT_files_for_mulea) and through the *muleaData* ExperimentData Bioconductor package (Ari *et al*., 2024) for 27 model organisms, from Bacteria to human (Supplementary Table 1). The ontologies are available in all major gene and protein identifiers (IDs), such as UniProt protein ID, Entrez, Gene Symbol, and Ensembl gene ID, for user convenience. Furthermore, *mulea* accepts the standardised GMT format, allowing easy integration of other data sources like MsigDB (Subramanian *et al*., 2005), Enrichr, *KEGG* (conversion script provided at the ELTEbioinformatics/GMT_files_for_mulea GitHub repository), or even user-defined ontologies.

Traditional enrichment analysis methods also suffer from the overcorrection of *p*-values. Specifically, conventional *p*-value adjustment methods, like the Bonferroni (1936) and Benjamini-Hochberg (1995), often overcorrect for multiple testing as they fail to account for the inherent interconnectedness among gene sets and ontology terms. Therefore in the *mulea* package, we introduce a re-sampling-based, *empirical false discovery rate* (eFDR) correction method.

## Methods and features in *mulea*

### The set-based *ORA* approach

*mulea* implements a set-based enrichment analysis approach that utilizes the hypergeometric test to identify statistically significant overrepresentation of elements from a query set (*e*.*g*., significantly upregulated genes) within a background set (*e*.*g*., all investigated genes). Therefore a predefined threshold value – such as 0.05 for the corrected *p*-values and *z*-scores, or 2-fold change – has to be used in the preceding analysis. To determine overrepresentation in ontology entries, *mulea* employs the hypergeometric test, which is analogous to the one-tailed Fisher’s exact test.

### Addressing multiple testing: *p*-value correction in the ORA analysis

Performing numerous statistical tests, such as evaluating enrichment across all ontology entries, leads to an inflated number of significant results (*p*-values < 0.05) due to chance, even if all null hypotheses are true. This phenomenon, known as the multiple testing problem, necessitates *p*-value correction. *mulea* offers various methods, including Bonferroni, Benjamini-Hochberg, and empirical false discovery rate (eFDR) correction.

However, Bonferroni and Benjamini-Hochberg methods assume independent tests, which rarely holds true in functional enrichment analyses. For example, GO categories exhibit a hierarchical structure, potentially leading to an unnecessary exclusion of significant results (enriched entries). Therefore, *mulea* implements the eFDR correction, which takes into account the distribution of test statistics, making it better suited for analyzing gene sets and ontologies typically employed by biologists. The eFDR implementation is based on methods described by Reiner *et al*. (2003) and Hastie *et al*. (2009). Detailed explanations of the eFDR algorithm and its advantages over the Benjamini-Hochberg method are provided in the Supplementary Notes.

It is important to note that eFDR correction can be computationally intensive, especially when dealing with large ontologies and numerous resampling rounds (we recommend at least 10,000). To address this, *mulea* implements the eFDR functionality in efficient C++ code. While similar approaches exist in tools like *Gowinda* (Kofler and Schlötterer, 2012) and *FuncAssociate* (Berriz *et al*., 2009), *mulea* offers advantages in terms of data type compatibility and offline usability.

The ranked list-based GSEA approach *mulea* facilitates ranked list-based enrichment analysis using the GSEA approach. This method requires an ordered list of elements (*e*.*g*., genes) as input, where the order reflects the user’s prior analysis (*e*.*g*., based on *p*-values and/or fold-changes). The list should encompass all elements involved in the analysis, such as all expressed genes in a differential expression study. *mulea* leverages the Kolmogorov-Smirnov statistic coupled with a permutation test (Subramanian *et al*., 2005) to assess enrichment within gene sets. This implementation is achieved through integration with the *fgsea* package (Korotkevich *et al*., 2021) from Bioconductor.

### Refining enrichment analysis by filtering ontology entries

Enrichment analysis results can sometimes be skewed by overly specific or overly broad ontology entries. *mulea* empowers users to address this issue by enabling the exclusion of such entries from the analysis. This filtering capability allows researchers to ensure that the results better match the expected scope.

### Results presentation and visualization

*mulea* offers various formats for presenting enrichment analysis results. By default, both ORA and GSEA results are provided in a tabular format. Additionally, users can leverage the *mulea* package to generate diverse visualizations, including lollipop and barplot charts, networks, and heatmaps (Figure 1).

**Figure 1:**
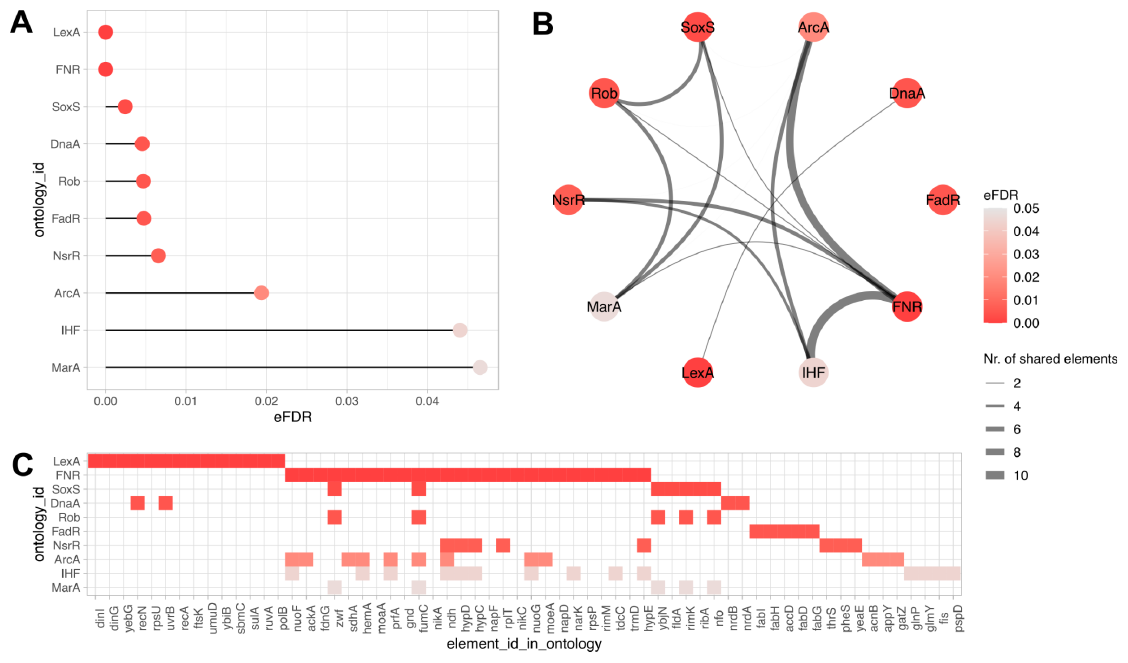
Visualization examples for an overrepresentation analysis. Visualization of overrepresented transcription factors among significantly upregulated *Escherichia coli* genes (GSE55662 Gene Expression Omnibus experiment), using the Regulon database (Salgado *et al*., 2023). Transcription factors regulating less than 3 or more than 400 target genes were excluded. (A) Lollipop chart: visualises the distribution of eFDR values (x-axis) < 0.05 for enriched transcription factors (y-axis). (B) Network representation: nodes represent enriched transcription factors (coloured based on eFDR values), while edges connect nodes sharing at least one target gene among the significantly upregulated genes, and are weighted by the number of such shared genes. (C) Heatmap: illustrates which elements (target genes, *x*-axis) belong to enriched ontologies (transcription factors, *y*-axis). Cell colours correspond to eFDR values.

## Comparison and Discussion

Here we present the *mulea* R package, offering a unique combination of features for functional enrichment analysis. *mulea* integrates two enrichment approaches (ORA and GSEA) with an empirical false discovery rate (eFDR) correction method, providing robust statistical assessments. Additionally, *mulea* encompasses diverse ontologies for enrichment analysis across multiple species, data types, and identifiers, catering to a broad range of research needs. While some functionalities overlap with existing software (Supplementary Table 2), *mulea* presents a comprehensive solution, uniting advanced methods and gene sets within a single package. This streamlined approach simplifies the analysis process and facilitates the interpretation of high-throughput results. While *mulea* shares functional similarities with tools like *Enrichr, g:Profiler*, and *clusterProfiler*, it offers several distinct advantages. Notably, compared to these tools, *mulea* implements a more rigorous multiple-testing correction method (eFDR) making the analysis more sensitive to detect significant enrichments. *mulea* provides pre-defined gene sets for a broad range of organisms by including 27 species. Thus, *mulea* extends beyond an established gene set collection, MSigDB, which is limited to human and mouse. This broader species coverage enhances the applicability of *mulea* to various research contexts. Furthermore, *mulea* empowers users to incorporate their own ontologies using a dedicated function, enabling them to leverage datasets from diverse sources and extend the analytical scope beyond the default options. These unique features establish *mulea* as a versatile and user-friendly resource for researchers conducting functional enrichment analyses.

## Acknowledgements

We thank Tamás Korcsmáros for his helpful comments. We dedicate this package to Péter Forgács, whose early contributions to developing the code laid the groundwork for this tool.

