## Supplementary Note for "mulea - an R package for enrichment analysis using multiple ontologies and empirical FDR correction"

for

### The ontology files provided to the *mulea R* package

#### Gene Matrix Transposed (GMT) format

The GMT format stores collections of genes or proteins associated with specific ontology terms. It uses a tab-delimited text file with one row per term. Each row consists of three key elements:

1. Ontology identifier: This column uniquely identifies the element within the referenced ontology.
2. Ontology name or description: This column provides a user-friendly label or textual description for the term.
3. Associated genes/proteins: This column lists identifiers (separated by spaces) of genes or proteins belonging to the term.

Within the *mulea*, these entities are named `ontology_id`, `ontology_name`, and `list_of_values`, respectively. Additionally, rows start with a "#" comment containing supplementary information about the referenced ontology, such as its type, source, species, version, and identifier.

##### Pre-defined GMT files

We provide GMT files generated from multiple different types of ontologies to make them appropriate for enrichment analyses. The GMT files are available for 27 species and are provided with multiple gene and protein identifiers (IDs). We recommend using the "primary identifiers" (listed in Supplementary Table 1) for each ontology if possible. Due to inequalities in identifier coverage, the *mulea* results may be slightly different when using alternative IDs.

##### Identifier mapping

UniProt's API (https://www.uniprot.org/help/id_mapping) facilitated the mapping of different ID types. We utilized the entire UniProt database (Swiss-Prot and TrEMBL) to map IDs to UniProt, Ensembl, Entrez (Gene ID), Gene symbol formats across all species, and Locus IDs where applicable. In some cases, species-specific IDs like FlybaseID (fruit fly) are included.

##### Availability

The GMT files are available at https://github.com/ELTEbioinformatics/GMT_files_for_mulea and can be directly read into the R project using the `read_gmt` function with the link of the file, for example:

```
mulea::read_gmt(file =
"https://raw.githubusercontent.com/ELTEbioinformatics/GMT_files_fo
r_mulea/main/Arabidopsis_thaliana_3702/Pathways_Reactome_Arabidops
is_thaliana_EnsemblID.gmt")
```

Or one can apply the *muleaData* ExperimentData Bioconductor package, see below.

#### The *muleaData* ExperimentData Bioconductor package

The *muleaData* Bioconductor R package, an ExperimentData package, simplifies access to these ontologies. It allows the user to download and utilize the ontologies as *R* objects in the familiar GMT data frame format using the ExperimentHub library. For example:

```
# Calling the ExperimentHub library.
library(ExperimentHub)

# Downloading the metadata from ExperimentHub.
eh <- ExperimentHub()

# Creating the muleaData variable.
muleaData <- query(eh, "muleaData")

# Checking the muleaData variable.
muleaData

# Looking for the ExperimentalHub ID of i.e. target genes of
transcription
# factors from TFLink in Caenorhabditis elegans.
mcols(muleaData) %>%
  as.data.frame() %>%
  dplyr::filter(species == "Caenorhabditis elegans" &
                sourceurl == "https://tflink.net/")

# Creating a variable for the GMT data.frame of EH8735.
```

```
# EH8735 contains small-scale measurement results, where the
target genes are
# coded with Ensembl ID-s
Transcription_factor_TFLink_Caenorhabditis_elegans_SS_EnsemblID <-
muleaData[["EH8735"]]
```

### Addressing multiple testing challenges: The eFDR correction

Traditional multiple testing corrections like *Bonferroni* and *Benjamini-Hochberg* assume independent tests, which rarely hold true in overrepresentation analysis due to inherent relationships between terms and gene sets. This can lead to overly conservative results and the loss of significant findings.

To address this limitation, *mulea* implements a more suitable re-sampling-based *empirical false discovery rate* (eFDR) correction method, that accounts for the test statistics distribution. Note that this procedure makes no reference to the original *p*-values at all, but rather works directly with the test statistics. eFDR correction effectively accounts for the interconnectedness within the data, leading to more accurate control of false discoveries and a better reflection of true positives in overrepresentation analysis results.

The specific eFDR correction method implemented in *mulea* aligns with the one used in the *Gowinda* software (Kofler and Schlötterer, 2012), as described in Reiner *et al.* (2003) and Hastie *et al.* (2009).

#### Calculating the eFDR

Let $I$ be the size of the ontology (*i.e.* the number of categories ($i$) in the ontology, *e.g.* the number of GO terms (The Gene Ontology Consortium, 2021)):

$$i=1,2,\ldots,I$$

$S$ the number of resampling steps ($s$):

$$s=1,2,\ldots,S$$

Then an observed ($R_{obs}$) and an expected ($R_{exp}$) count of element sets having *p*-values less than or equal to a threshold (where $p$ is the *p*-value that should be corrected) can be computed. The $eFDR$ can subsequently simply be calculated by dividing $R_{obs}$ by $R_{exp}$.

Where $R_{obs}$ is the number of cases where the actual $p$-value ($p_i$) is less than or equal to the $p$-values ($P$) of every ontology category ($O_i$):

$$R_{obs} = \sum_{i=1}^{I} \left(p_i \leq P\right)$$

$R_{exp}^{s}$ is the number of cases where the $p$-value ($p_i^s$) of the selected element set (same size as the investigated element set, randomly chosen from the background) is less than or equal to the $p$-values ($P$) of the investigated element set for every $O_i$:

$$R_{exp}^{s} = \sum_{i=1}^{I} \left(p_i^s \leq P\right)$$

Where $R_{exp}$ is the sum of $R_{exp}^{s}$-s for all $s$-s (resampling rounds) divided by $S$ (the number of resampling rounds):

$$R_{exp} = \frac{\sum_{s=1}^{S} R_{exp}^{s}}{S}$$

$$eFDR = \frac{R_{exp}}{R_{obs}}$$

Depending on the ontology size and the number of resampling rounds (min. 10,000 is suggested) the $eFDR$ correction can be computationally intense.

### Testing the *p*-value correction methods

To illustrate the differences between the *empirical false discovery rate* (*eFDR*) and *Benjamini-Hochberg p*-value correction methods, we conducted a simulation study. We performed overrepresentation analysis 9 million times (9000 iterations x 1000 repeats per iteration) using various noise levels. To perform the test we used an established ontology: the transcription factor and target gene interactions of *Homo sapiens*, measured with small-scale methods, implemented with uniprotIDs, as it can be found on the TFLink gateway (Liska *et al.*, 2022). We chose 20% of the transcription factors (terms of the GMT file) randomly and took 85% of the proteins belonging to each (elements of each term) as the overrepresented sets. We also added randomly chosen other proteins to each set representing the noise in biological data. To do so, different noise ratios were applied: 0, 0.1, 0.2, 0.3, 0.4, 0.5, 0.6, 0.7, and 0.8. Then we calculated the overrepresentation analysis 1000 times for each noise ratio using all the genes in the GMT file as background gene set and repeat the re-sampling for the eFRD p-value correction 1000 times. After that, we calculated the true and the false positive rates for each result (Supplementary Figure 1).

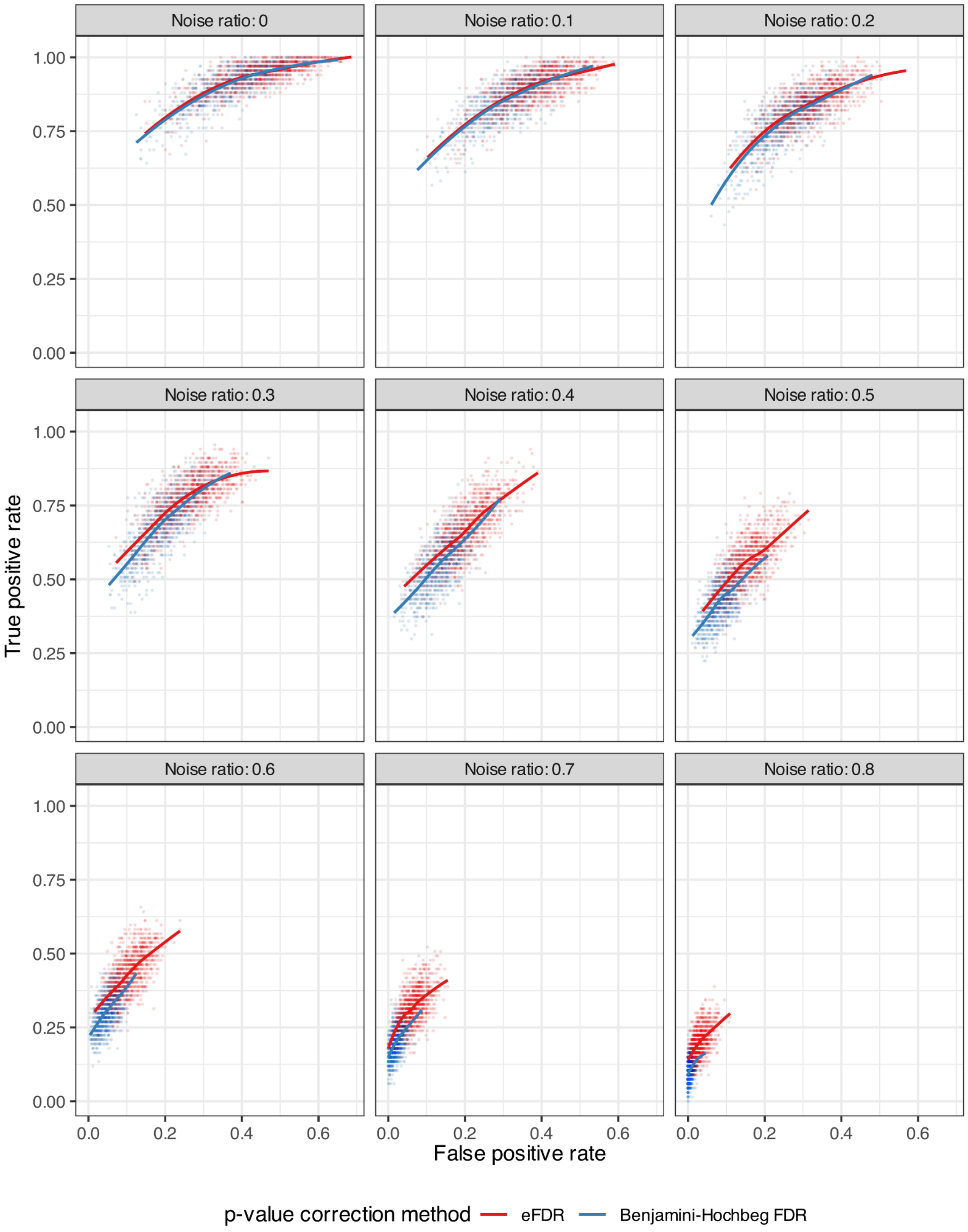


**Supplementary Figure 1: True and false positive rates of tests where the p-values were corrected using the empirical false discovery rate (eFDR) and the Benjamini-Hochberg methods.** Scatter plots of the true positive rates (y-axis) and false positive rates (x-axis) faceted by different noise ratios (from 0 to 0.8). The dots represent the true and false positive rates of individual experiments, while the lines are the results of local polynomial regression fittings. The dots and lines are coloured red when the eFDR p-value correction method was applied and blue when the Benjamini-Hochberg method was applied.

Using the *eFDR* method we always got significantly higher rates of true positives compared to the results with the *Benjamini-Hochberg* correction. The *p*-values of the one-sided paired *t*-test were less than 2.2e-16 for all the nine different noise ratios that were applied, which means that the *eFDR* correction method fits the actual data better than the *Benjamini-Hochberg* adjustment.
